# Rapid stress hardening in the Antarctic midge improves male fertility by increasing courtship success and preventing the decline of accessory gland proteins following cold exposure

**DOI:** 10.1101/2021.02.27.432016

**Authors:** Oluwaseun M. Ajayi, J. D. Gantz, Geoffrey Finch, Richard E. Lee, David L. Denlinger, Joshua B. Benoit

## Abstract

Rapid hardening is a process that quickly improves an animal’s performance following exposure to a potentially damaging stress. Features of reproduction can be improved by rapid hardening, but little is known about how rapid hardening may contribute to physiological responses in the cold environment of Antarctica. In this study of the Antarctic midge, *Belgica antarctica* (Diptera, Chironomidae), we examine how rapid hardening in response to dehydration (RDH) or cold (RCH) improves male pre- and post-copulatory function related to fertility when the insects are subsequently subjected to a damaging cold exposure. Neither RDH nor RCH improved survival in response to lethal cold stress, but male activity following sublethal cold exposure was enhanced. Both RCH and RDH improved mating success of males compared to those subjected directly to a sublethal bout of cold. Egg viability decreased following direct exposure to sublethal cold, but improved following RCH and RDH. Sublethal cold exposure reduced expression of four accessory gland proteins, while expression remained high in males exposed to RCH. Though rapid hardening may be cryptic in males, this study shows that it can be revealed by pre- and post-copulatory interactions with females.

## Introduction

The Antarctic midge, *Belgica antarctica*, is the only insect endemic to maritime Antarctica (Convey et al., 1996; Sugg et al., 1983). It has a sporadic distribution along the western coast of the Antarctic Peninsula and the South Shetland Islands, where the wingless adults may form aggregations of thousands under favorable conditions (Convey et al., 1996; Potts et al., 2020; Sugg et al., 1983). Larval development extends over two years; developmental progression occurs during the austral summer, and larvae overwinter frozen or immobile within the ice (Usher and Edwards, 1984). Females produce a single batch of eggs or sometimes two smaller clutches (Finch et al., 2020; Harada et al., 2014). The eggs are deposited within a gel matrix produced by the female accessory gland (Finch et al., 2020), and numerous proteins from the accessory gland are critical for nourishing developing larvae, preventing egg/larval dehydration, and thermally buffering the developing larvae. These proteins are produced during both the larval and adult stages and are synthesized by multiple organs. Consequently, stress exposure during larval stages can reduce reproductive output of the females (Finch et al., 2020).

During the austral summer, mating occurs in swarms where numerous males court and attempt to mate (Convey et al., 1996; Finch et al., 2020; Sugg et al., 1983). Males produce a suite of accessory proteins that are transferred to the females during mating, and based on sequence similarity to accessory gland proteins from other insects, these proteins are likely to impact female fertility and male sperm viability along with competition (Avila et al., 2011; Chapman et al., 2000; Simmons, 2019; Sitnik et al., 2016; Wolfner, 1997). As with females, stress exposure of male larvae impacts subsequent male fertility (Finch et al., 2020). It is not known how cold exposure impacts fertility when males are exposed to cold as adults. Impacts could be revealed during pre-copulation competition between males or post-copulation by a reduction in accessory gland protein or viable sperm. Cold exposure reduces male fertility in multiple insect systems (Chakir et al., 2002; Lacoume et al., 2007; Rinehart et al., 2000; Singh et al., 2015; Vollmer et al., 2004).

Rapid cold hardening (RCH) is a highly-responsive protective process documented in many insect systems against cold injury (Teets et al., 2019; Teets et al., 2020). RCH can also be induced by short periods of dehydration, which we refer to as rapid dehydration hardening (RDH) (Levis et al., 2012). Mechanisms underlying processes of RCH and prolonged cold acclimation include synthesis and accumulation of specific cryoprotectants, changes in the composition of cell membranes, inhibition of apoptotic pathways, changes in stress protein abundance, protein phosphorylation shifts, and metabolic differences (reviewed in (Teets and Denlinger, 2013; Teets et al., 2020). Importantly, RCH and cold acclimation offer beneficial effects allowing biological processes such as movement, feeding, and mating to occur at lower temperatures or to enhance performance when conditions become more favorable (Kelty et al., 1996; Shreve et al., 2004; Srithiphaphirom et al., 2019; Teets et al., 2019). During summer, males of the Antarctic midge are unlikely to be exposed to lethal temperatures (-8 to -10°C) because summer temperatures in their habitat commonly range from -5 to 5°C. In summer-acclimated larvae, RCH improves recovery performance following sub-lethal freezing injury and facilitates energy saving (Teets et al., 2019). The impact of RCH on adult reproductive performance has yet to be examined even though adults are active and moving on the surfaces of the buffered habitats where the larvae reside.

In this study, we examined the impact of RCH and RDH on male performance of *B. antarctica* following exposure to sublethal cold stress. Specifically, we assessed pre- and post-copulatory function related to male performance and fertility. Males exposed directly to sublethal cold stress suffered significant declines in mating competitions with untreated males and subsequent fertility was reduced. These impairments were reduced by RCH or RDH prior to mating. The RCH-induced enhancement of male competitiveness is likely due to a general increase in activity, and the boost in fertility is likely due to retention of the capacity to express seminal fluid proteins. These physiological responses suggest that cold hardening may be essential for allowing males to retain fertility during the austral summer.

## Materials and Methods

### Midge collection

Adults and larvae were collected from the same location, on off-shore islands near Palmer Station, Antarctica. Larvae within organic debris were returned to Palmer Station and extracted with a modified Berlese funnel (Benoit et al., 2007b). Following recovery, larvae were stored with substrate from their natural habitat (rocks, soils, moss, and the alga *Prasiola crispa*, which serves as a food source for *B. antarctica*) at 2-4°C. Larvae were shipped to the University of Cincinnati and stored under similar conditions until they emerged as adults and were used in the studies described. Males and females were separated based on their morphological characters.

### Survival assessment

Survival following cold exposure was based on previous methods (Benoit et al., 2007a; Lee et al., 2006) but adapted for these specific experiments. Adult males held at 4°C were placed into individual 2 ml microcentrifuge tubes and transferred from 4°C to either -5°C for 1 h (sublethal cold exposure) or - 10°C for 1 h (lethal cold exposure), based on temperatures determined in previous studies (Lee et al., 2006). RCH was induced by exposure to -2°C for 1 h before direct transfer to the treatment temperature. RDH was accomplished by exposing adults to 75% RH until a loss of 5-8% of their water content (Benoit et al., 2007a; Benoit et al., 2009) before cold exposure. Survival was assessed following a 24 h recovery at 4°C. Adults were deemed alive if the individual could move at least 5 cm. Three groups with 8 replicates were examined for each treatment. A one-way ANOVA was used to examine significance and a Tukey’s test was used to identify differences between treatments.

### Activity

A Locomotor Activity Monitor (TriKinetics Inc., Waltham, MA, USA) in conjunction with DAMSystem3 Data Collection Software (TriKinetics) was used to assess activity of adult males. Individuals were placed in standard *Drosophila* vials (25 mm diameter and 95 mm height) following 6 h of recovery at colony conditions of 4°C. Tubes were placed horizontally in the monitor and the entire system was placed within a plastic container held at 93% RH using a saturated salt solution of potassium nitrate. Temperature was held at 4°C within an environmental chamber. After a two-hour acclimation, general activity was measured for 4 h under continuous light. For each treatment, 15-20 males were monitored. A one-way ANOVA was used to examine significance and a Tukey’s test was used to identify differences between treatments.

### Male mating competition

Competitive mating assays were conducted following cold exposure or rapid hardening followed by cold exposure. Cold level for exposure was sublethal (-5°C for 1 h). RCH was induced by exposure to -2°C for 1 h before direct transfer to the treatment temperature. Following treatment, males were returned to colony conditions and allowed to recover for 6 h prior to mating assays to match studies on activity. The mating assays were accomplished by releasing a virgin female into an arena with two males. Males were distinguished by partial removal of either the left or right antennae (randomized between each assay). The first male to copulate with the female was scored as a successful copulation. Each assay was replicated 50 times and significance was determined with a Chi-squared (χ^2^) test. New females and males were used for each assay.

### Fertility assay

To determine the impact of thermal exposure on male fertility, males exposed to thermal stress were allowed to mate with either two females based on previously developed methods (Finch et al., 2020). Briefly, virgin females were collected immediately after adult eclosion and stored at 4°Cand 93% RH, with access to moist substrate. Males were exposed to thermal stresses as described in the survival assessment section. Following treatment, males were evaluated for survival and allowed to mate with two females consecutively. Following mating, females were returned to colony conditions of 4°Cand 93% RH and observed for deposition of eggs. Eggs, held at 4°Cand 93% RH, remained attached to the wet substrate and were observed for larval emergence for 60 days (Finch et al., 2020; Harada et al., 2014). Each treatment was replicated with 8-12 males. A one-way ANOVA was used to examine significance followed by a Tukey’s test for differences between treatments.

### RNA-seq analysis to identify highly enriched male accessory glands genes for targeted studies

To establish putative underlying factors that could impact fertility, we re-examined RNA-seq data on male and male accessory glands (Finch et al., 2020), with the goal of identifying four targets for subsequent analyses following cold exposure and RCH. The desired targets were selected on the basis of having high expression in males and male accessory glands compared to females/larvae and female accessory glands. The four targets were characterized by BLAST comparison to other genes in other insect systems. The four selected targets included: IU25_04442, a *Belgica* specific gene; IU25_01011, a seminal metalloprotease; IU25_12390, an antigen 5-like allergen; IU25_12518, a seminal metalloprotease.

### qPCR analyses of male accessory proteins

RNA was extracted from the accessory glands by homogenization (BeadBlaster 24, Benchmark Scientific) in Trizol reagent (Invitrogen), using manufacturer’s protocols with slight modification based on other studies of invertebrates. Extracted RNA was treated with DNase I (Thermo Scientific) and cleaned with a GeneJet RNA Cleanup and Concentration Micro Kit (Thermo Scientific) according to the manufacturer’s protocols. RNA concentration and quality were examined with a NanoDrop 2000 (Thermo Scientific).

qPCR analyses were conducted based on previously developed methods (Finch et al., 2020; Hagan et al., 2018; Meibers et al., 2019). RNA was extracted as described previously for independent biological replicates. Complementary DNA (cDNA) was generated with a DyNAmo cDNA Synthesis Kit (Thermo Scientific). Each reaction used 250 ng RNA, 50 ng oligo (dT) primers, reaction buffer containing dNTPs and 5 mmol•l^−1^ MgCl_2_, and M-MuLV RNase H+ reverse transcriptase. KiCqStart SYBR Green qPCR ReadyMix (Sigma Aldrich, St Louis, MO, USA) along with 300 nmol l^−1^ forward and reverse primers, cDNA diluted 1:20, and nuclease-free water were used for all reactions. Primers were designed using Primer3 based on contigs obtained from the transcriptome analysis (Table S1). qPCR reactions were conducted using an Illumina Eco quantitative PCR system. Reactions were run according to previous studies (Finch et al., 2020; Hagan et al., 2018; Meibers et al., 2019). Three biological replicates were examined for each sex, and two biological replicates were examined for each accessory gland. Expression levels were normalized to *rpl19* using the ΔΔCq (Finch et al., 2020; Teets et al., 2013). A one-way ANOVA was used to examine significance followed by a Tukey’s test for differences between treatments.

## Results

### Rapid hardening does not impact survival of males

Significant differences in mean survival were noted for the different treatment groups as determined by one-way ANOVA (*F*_8,18_ = 34.25, *p-value* = 2.38 × 10^-9^). Sudden exposure of males to the extreme cold treatment significantly decreased survival in comparison with controls, but survival was not improved by RCH (Fig. 1). Although the value for male survival was higher when extreme cold exposure was preceded by RDH, this increase was not significant (Fig. 1). Neither RCH nor RDH improved survival in those exposed to a sublethal stress (Fig. 1). Thus, we observed no evidence that rapid hardening enhanced survival of male adults.

**Figure 1.**
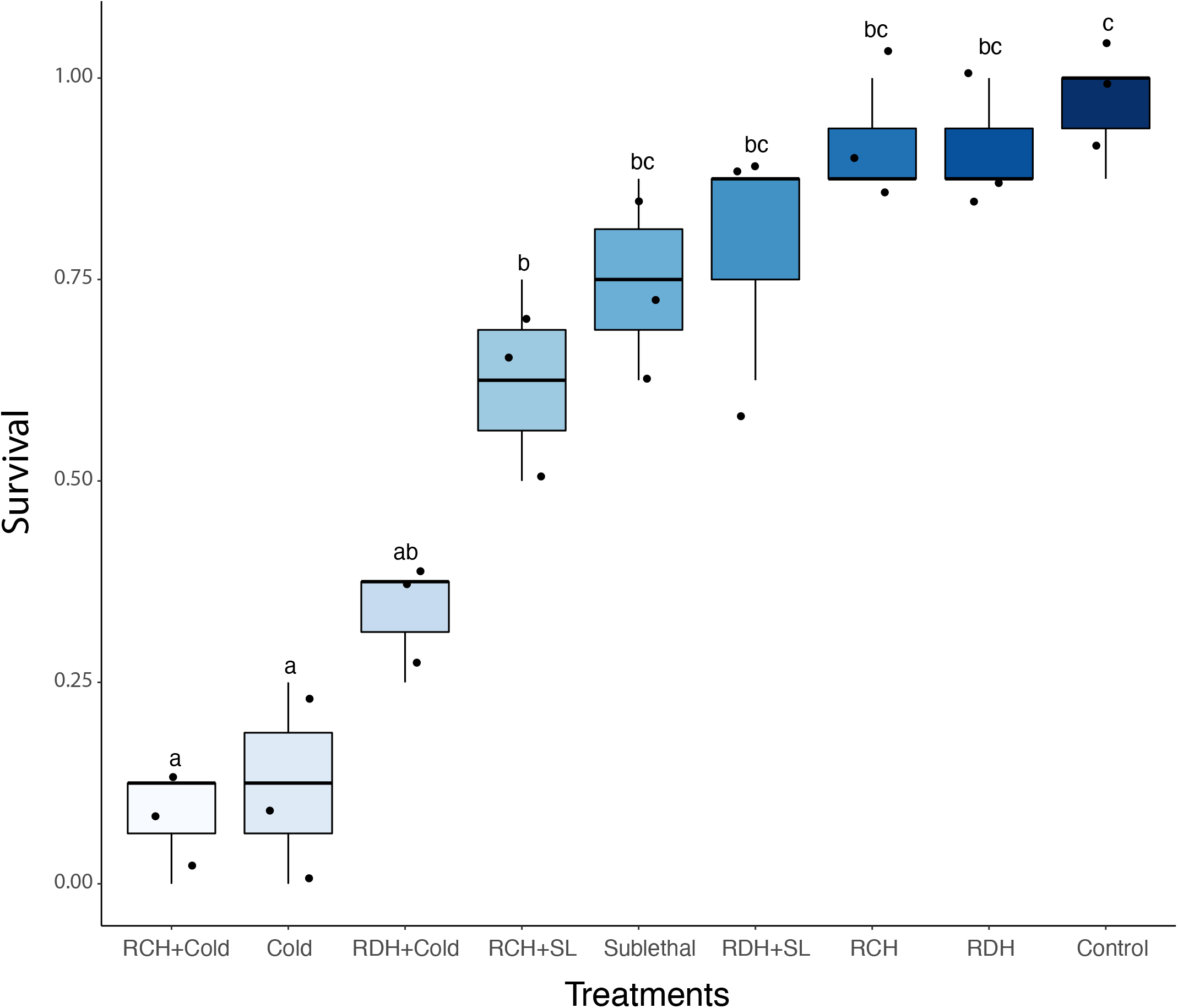
Effect of rapid hardening on male survival of the Antarctic midge. Different letters indicate significant (one-way ANOVA, Tukey’s Honest Significant Difference test, *P-value* < 0.05) differences between treatment groups. (RCH+Cold = rapid cold hardening before damaging cold exposure; Cold = damaging cold exposure; RDH+Cold = rapid dehydration hardening before damaging cold exposure; RCH+SL = rapid cold hardening before sublethal cold stress; Sublethal = sublethal cold stress; RDH+SL = rapid dehydration hardening before sublethal cold stress; RCH = rapid cold hardening; RDH = rapid dehydration hardening, Control = untreated group).

### Rapid hardening improves male activity and competition during female courtship

Sublethal cold exposure resulted in significant reduction in activity (*F*_3,72_ = 12.07, *p-value* = 1.73 × 10^-6^), but activity was retained in RCH-treated compared to control males (Tukey’s test, *p-value* = 0.246, Fig. 2a). Activity was not significantly different between control males and those that experienced RCH. These results indicate that RCH allows males to maintain activity following cold stress.

**Figure 2.**
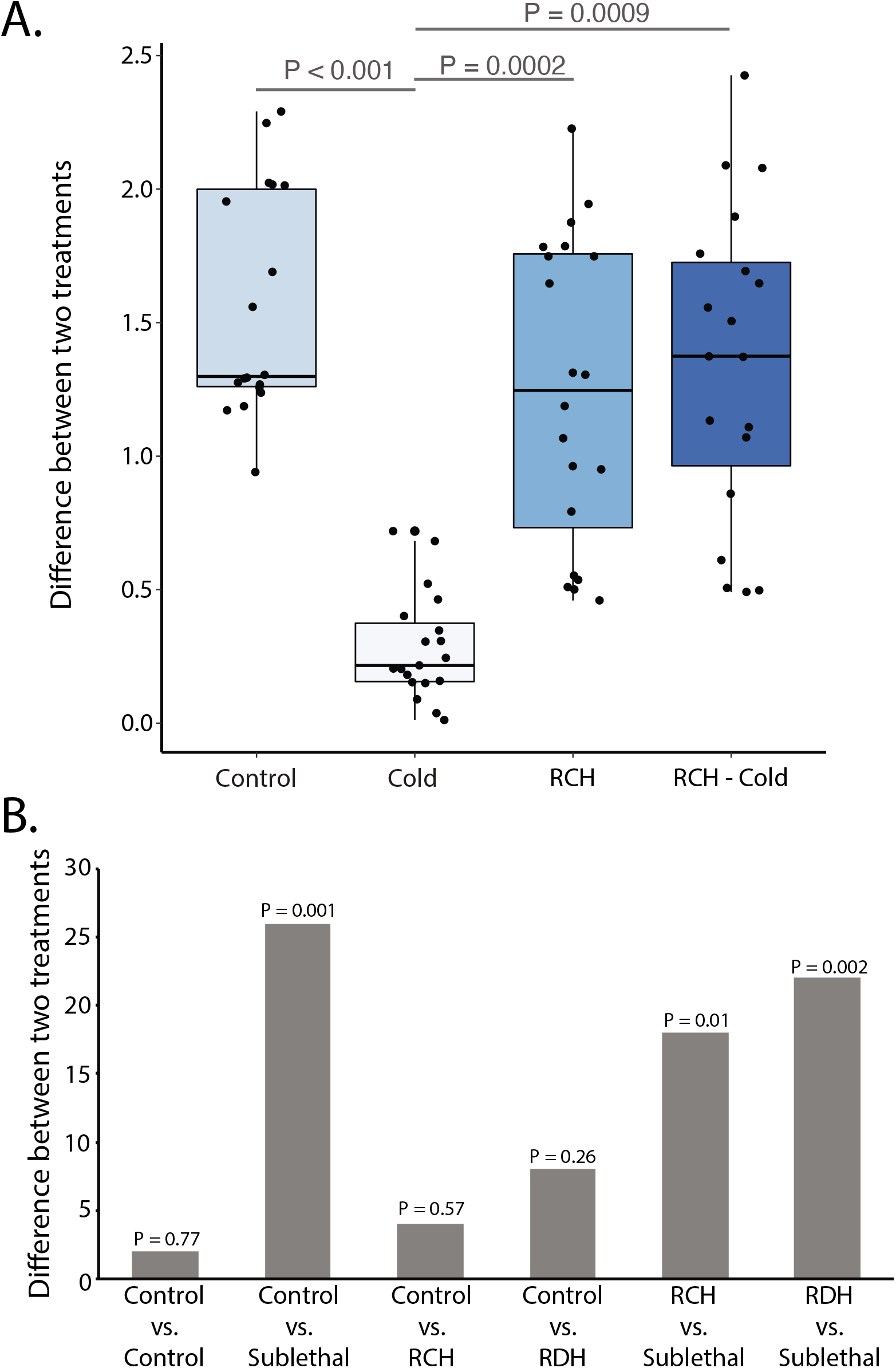
Effect of rapid hardening on competition between males of the Antarctic midge during courtship. (A) Activity differences in control, sublethal cold exposure, and rapid cold hardening (RCH) followed by sublethal cold exposure. Significance determined by ANOVA followed by a Tukey’s test for differences between treatments. (B) Difference between the choice of control or RCH males compared to the specific treatment. Significance was determined with χ^2^ test.

Results from the competitive mating assays demonstrated that sublethal cold exposure drastically impaired male mating (*N* = 50, χ^2^, *p-value* = 8.16 × 10^-5^, Fig. 2b). RCH or RDH without sublethal cold stress did not alter male mating (*N* = 50, χ^2^, *p-value* > 0.26, Fig. 2b). Males first exposed to RDH or RCH before sublethal cold stress showed increased mating compared to those only experiencing sublethal cold stress (*N* = 50, χ^2^, *p-value* < 0.01, Fig. 2b). These results suggest that RDH and RCH will likely improve pre-copulatory interactions in males, allowing for increased mating success.

### Rapid hardening prevents reductions in male fertility

The total number of eggs produced did not vary among treatment groups (one-way ANOVA, *p-value* = 0.456). But, egg viability significantly differed among treatments (Fig. 3.; one-way ANOVA, *F*_3,46_ = 14.12, *p-value* = 1.18 × 10^-6^). In particular, egg viability was significantly reduced (by nearly 40%) when eggs were exposed to sublethal cold (Tukey’s test, *p-value* < 0.001). This reduction in egg viability was prevented when males were pretreated with RCH (Fig. 3; *p-value* = 0.717). RCH before the sublethal cold exposure thus allowed males to maintain fertility, as indicated by egg viability (Fig. 3; *p-value* = 0.371 vs. control, *p-value* = 0.0009 vs. sublethal cold). Thus, both pre-copulatory and post-copulatory aspects of reproduction are impacted when males are exposed to sublethal cold stress.

**Figure 3.**
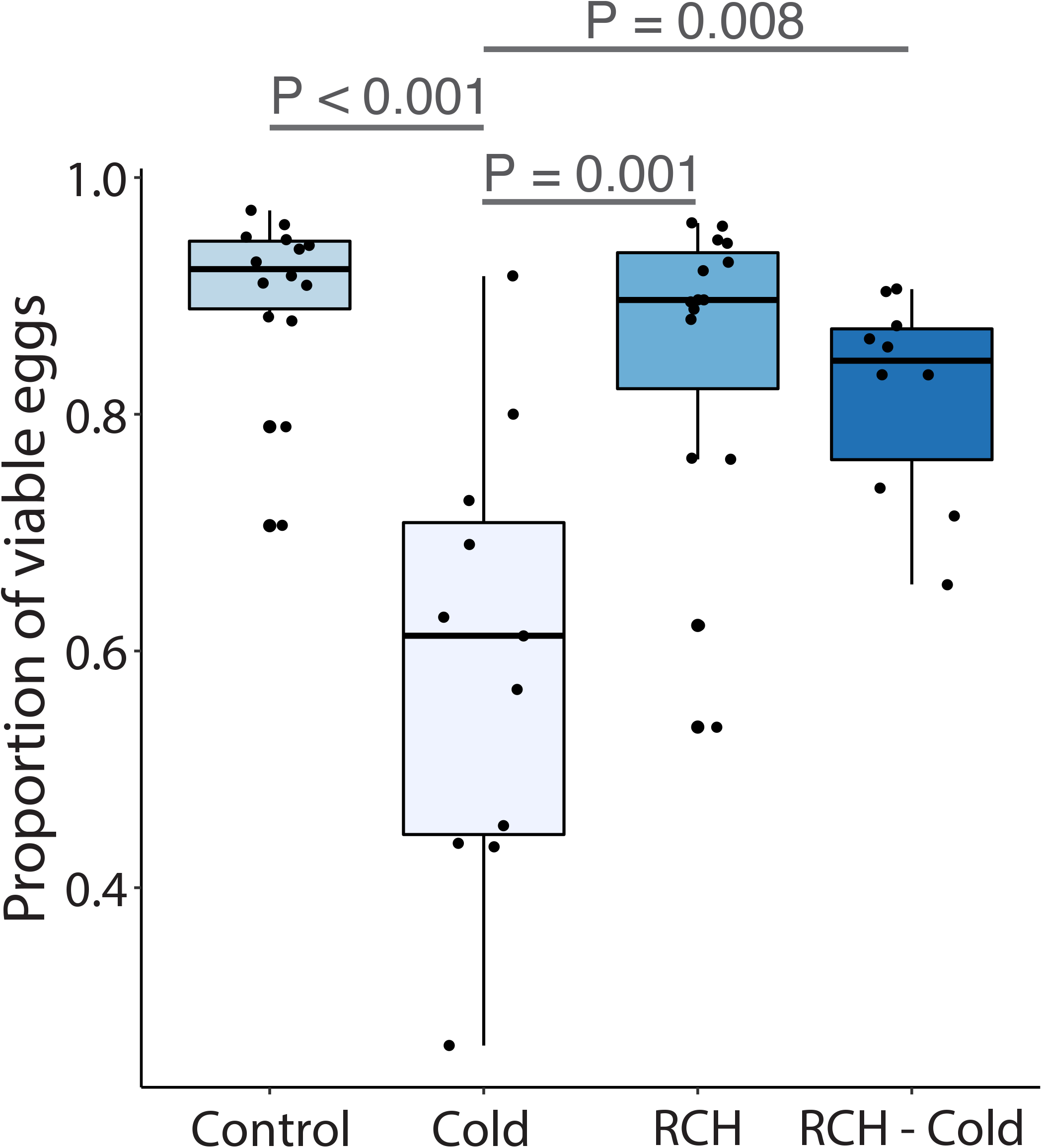
Effect of rapid hardening of males on viability of eggs produced by females. Treatments are control, cold exposure, rapid cold hardening (RCH), rapid cold hardening before sublethal cold exposure (RCH-Cold). Significant differences between treatment groups were determined with a one-way ANOVA followed by a Tukey’s test for differences between treatments. The total number of eggs produced by females did not vary between treatments (P = 0.456).

To determine a putative mechanism for reduced fertility, we examined expression patterns of four accessory gland proteins (ACPs) that are expressed exclusively in the male accessory gland (Fig. 4a). When exposed to sublethal cold stress, expression of all four ACPs was significantly reduced (Fig. 4b, one-way ANOVA, *p-value* < 0.03 in all cases). When subjected to RCH prior to sublethal cold exposure, expression of three ACPs increased in comparison to expression levels seen following sublethal cold exposure without RCH (Fig. 4b). These results suggest that a reduction in factors associated with seminal fluid production could underlie the observed reduction in male fertility.

**Figure 4.**
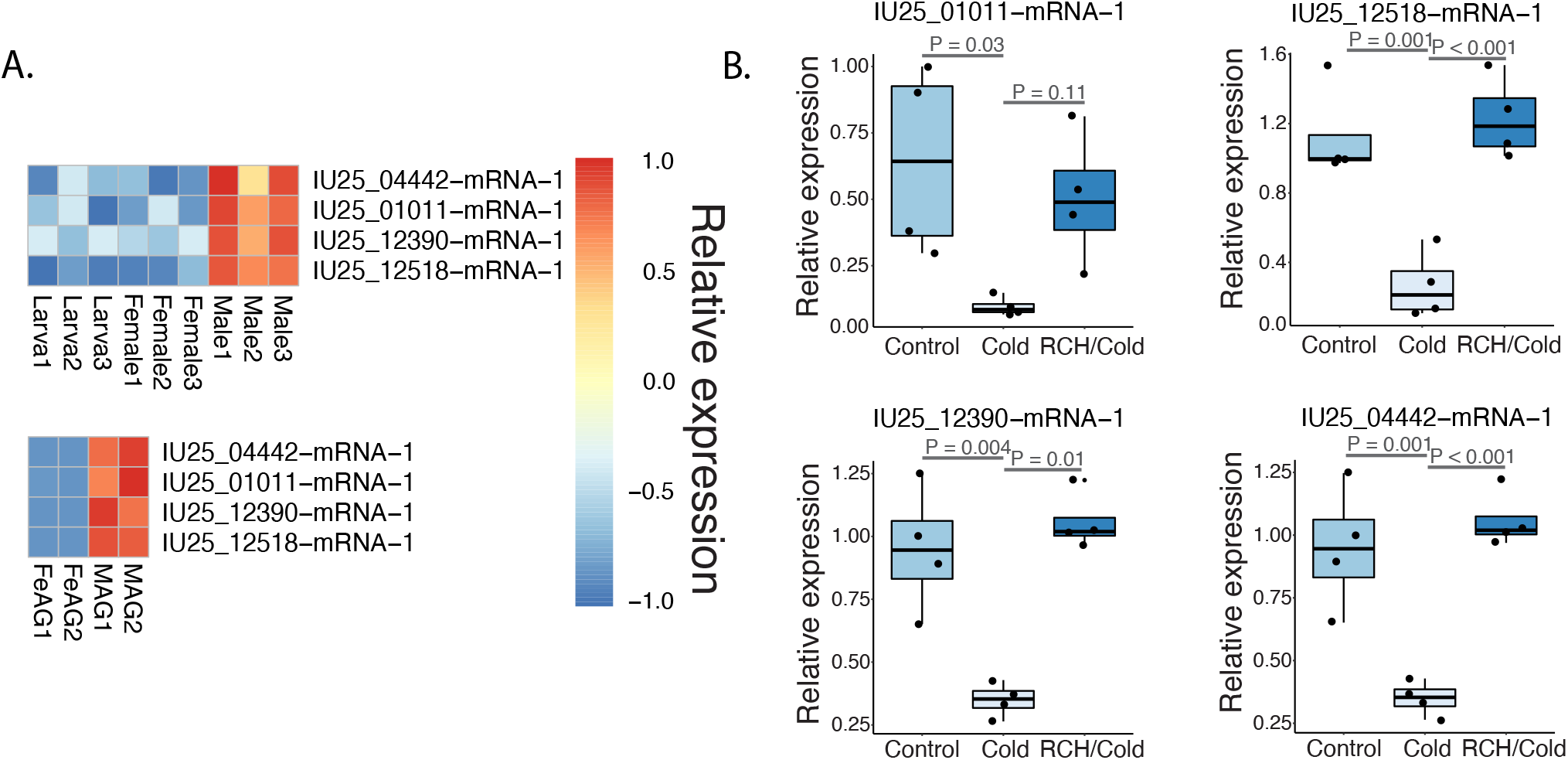
Transcript level analysis of male accessory gland proteins during cold exposure. (A) Expressional analysis to identify male accessory gland protein targets based on previous RNA-seq studies underlying reproductive biology of the Antarctic midge (Finch et al., 2020). (B) quantitative PCR analysis following cold exposure and rapid cold hardening (RCH). Significant differences between treatment groups were determined with a one-way ANOVA followed by a Tukey’s test for differences between treatments.

## Discussion

This study highlights the critical role of rapid hardening in relation to male performance in the Antarctic midge, as summarized in Figure 5. Hardening had very little impact on survival of males but allowed them to retain higher levels of activity. This increased activity is likely a major underlying factor allowing males to be more competitive in their ability to copulate with females. Male exposure to sublethal cold stress resulted in a significant reduction in fertilization rate, as reflected in production of fewer viable eggs. This impairment was recovered when males were subjected to RCH prior to cold exposure. This reduction in fertility could be due, in part, to a major reduction in expression of specific accessory gland proteins, which are critical for males to maintain maximum fecundity. The combined impact is that rapid hardening can promote male fitness in two ways: 1) by allowing high performance during pre-copulatory interactions, and 2) by allowing males to maximize fertility. Interactions between pre- and post-copulatory factors work together in contributing to male fertility (Birkhead and Pizzari, 2002; Polak et al., 2021). In this study, we also show that both aspects can be negatively impacted by exposure to stress.

**Figure 5.**
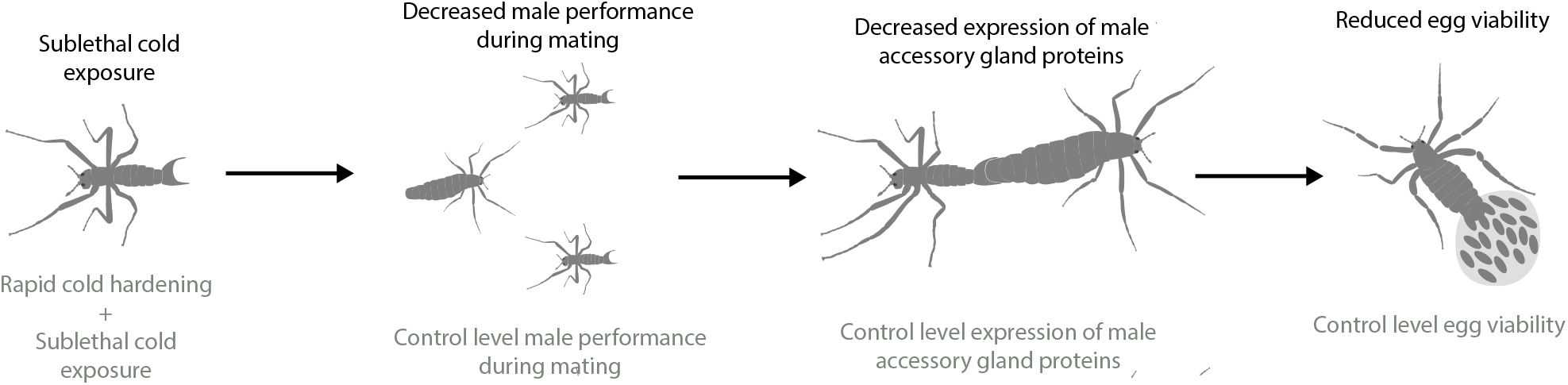
Summary of the impact of sublethal cold exposure and rapid cold hardening on pre-copulatory and post-copulatory features of male fertility and performance.

Increased survival during lethal cold stress is a major factor associated with rapid hardening, induced by dehydration or cold (Benoit et al., 2009; Teets et al., 2019; Teets et al., 2020). For the Antarctic midge, RCH significantly improves larval survival when exposed to freezing temperatures (Lee et al., 2006; Teets et al., 2008). But, the Increased survival appears to be stage-specific in the Antarctic midge. While larvae show improved survival following RCH, adults do not (Lee et al. 2006). Our results are consistent with these earlier observations showing that rapid hardening has little impact on adult survival of the midge. Rapid hardening in other insect systems, such as *Drosophila*, also shows differences between developmental stages, specifically in relation to survival (Colinet and Hoffmann, 2012; Jensen et al., 2007; Teets et al., 2020; Terblanche et al., 2007). Increased survival generated by RCH may lack ecological relevance for these short-lived adult midges because they are rarely exposed to potentially lethal temperatures during their active period in the austral summer (Lee et al., 2006; Teets et al., 2019).

A key aspect of RCH is organismal performance following exposure to sublethal temperatures (reviewed in (Teets et al., 2020)). Three key aspects related to sublethal cold stress are a lowering of the critical thermal minimum temperature (CT_min_), increased recovery from the CT_min_, and fitness/energetic advantages following cold stress (Benoit et al., 2021; Coulson and Bale, 1992; Findsen et al., 2013; Rinehart et al., 2000; Shreve et al., 2004; Teets et al., 2019; Teets et al., 2020). We did not determine whether RCH reduces the CT_min_ or recovery, but rapid hardening did improve the male activity levels to near those of controls following sublethal cold exposure. RCH likely prevents protein and membrane damage including resting potential depolarization that would normally occur following sublethal cold stress (Koštál et al., 2004; Koštál et al., 2006; Overgaard and MacMillan, 2017). Damage due to sublethal stress is likely most pronounced in muscle and nervous tissues (Andersen et al., 2015; Koštál et al., 2006; MacMillan et al., 2014; Overgaard and MacMillan, 2017), which in turn can be expected to significantly impact general activity levels, as observed for males in our study.

The overall increased level of activity is likely a major factor underlying the RCH-generated improvement in male copulatory behavior. *B. antarctica* mates in swarms (Finch et al., 2020; Sugg et al., 1983), thus there is likely significant pre-copulatory competition between males, and improved male performance could represent a significant advantage if there are periods of unexpected cold during the summer. In *Drosophila melanogaster*, courtship behavior is preserved by RCH at low temperatures (Shreve et al., 2004), but whether this improves courtship abilities following more extreme stress is not known. In larvae of the Antarctic midge, RCH does improve performance when larval locomotion is monitored (Teets et al., 2019). Studies from multiple systems suggest that RCH improves general activity during cold exposure, especially in chill susceptible systems (Overgaard and MacMillan, 2017)

Along with a reduction in pre-copulatory performance, male fertility declined following cold exposure as evidenced by a nearly 40% reduction in egg viability. This reduction was prevented when males were exposed to RCH before the sublethal cold stress. When pharate adult flesh flies, *Sarcophaga crassipalpis*, are exposed to cold stress, there is also a reduction in male fertility that can similarly be recovered by RCH (Rinehart et al., 2000). Among potential mechanisms that could contribute to this impairment, we noted that expression of previously identified male ACPs (Finch et al., 2020) were reduced substantially. RCH yielded a substantial recovery in expression of transcripts encoding the ACPs. Two of these proteins were identified as putative metalloproteases previously implicated in processing of seminal proteins, sperm competition, and sperm storage (Avila and Wolfner, 2017; LaFlamme et al., 2012; Laflamme et al., 2014). Along with sperm viability, female sexual receptivity also increases when specific proteases are inhibited (LaFlamme et al., 2012). This could have a substantial impact in species such *as B. antarctica* where mating occurs in swarms and females mate more than once (Finch et al., 2020). The two other ACPs examined here include one that does not have an assigned function and one that is related to antigen 5-like allergens. Antigen-like proteins are common in salivary glands in many insect systems (King and Spangfort, 2000), but the specific role for this protein in the Antarctic midge remains unknown. From our studies, we cannot rule out direct damage to sperm as an underlying cause for females producing fewer viable eggs, as documented in other insect systems (Colinet and Hance, 2009; Levie et al., 2005; Rinehart et al., 2000). Both the reduction in transcript levels for ACPs and sperm damage may contribute to reproductive impairment by low temperature, but regardless of the immediate cause, RCH appears to be an effective mechanism for protecting not only pre-copulatory activity but also post-copulatory factors impacting fertility in the Antarctic midge.

## Acknowledgments

This work was supported by the National Science Foundation grant DEB-1654417 (partially) and the United States Department of Agriculture 2018-67013 (partially) to J.B.B. for reusable equipment, National Science Foundation grant OPP-1341393 to D.L.D., and National Science Foundation grant OPP-1341385 to R.E.L. We thank the staff at Palmer Station, Antarctica for assistance in logistics, experiments, and helping make the field season a pleasant experience.

## Figure and table legends

**Table S1.**
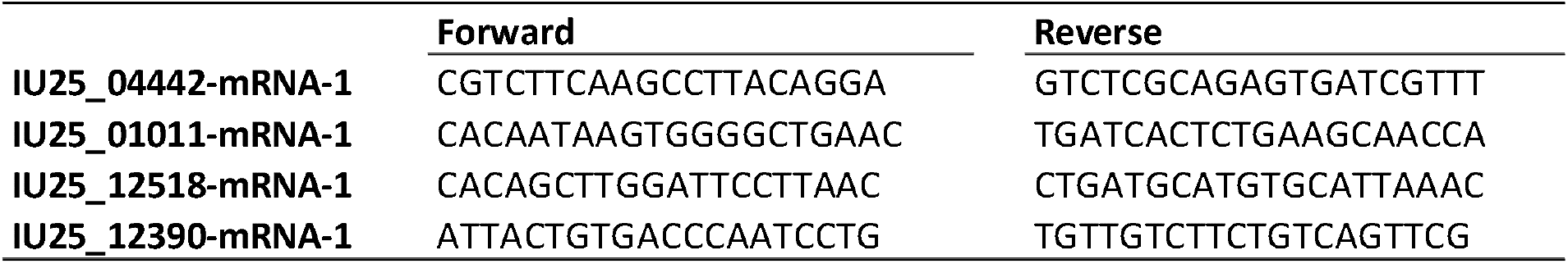
Primers utilized for quantitative PCR studies.

## Notes

### Competing Interest Statement

The authors have declared no competing interest.

